# Cholinergic and adrenergic modulation sculpts parallel inhibition of SST-INs in visual cortex

**DOI:** 10.64898/2026.10.28.762553

**Authors:** Jyoti Gupta, Chiayu Q. Chiu, Fabio Veneto, Julijana Gjorgjieva, Michael J. Higley

**Author notes:** **Correspondence:** Michael J. Higley, Department of Neuroscience, Yale University.

## Abstract

GABAergic interneurons are fundamental to the normal function of neuronal circuits in the neocortex. A growing literature has highlighted the role of somatostatin-expressing interneurons (SST-INs) in regulating cortical activity, and inhibition of SST-INs by other interneurons has emerged as a circuit motif leading to disinhibition of pyramidal neurons. However, the diversity and regulation of GABAergic inputs to SST-INs are only partially understood. Here, we combined widefield and 2-photon optogenetic stimulation of parvalbumin (PV)- and vasoactive intestinal peptide (VIP)-expressing interneurons with recordings from fluorescently identified SST-INs in layer 2/3 of primary visual cortex (V1). Our results show that both PV- and VIP-INs form inhibitory inputs onto SST-INs but with varying functional properties. PV-INs evoke strong, fast inhibition onto perisomatic regions of their target cells, while VIP-INs evoke slower inhibition onto target cell dendrites that is partially mediated by α5 subunit-containing GABA receptors. Cholinergic modulation through muscarinic receptors strongly suppresses PV-IN output while adrenergic modulation through beta receptors enhances VIP-IN output. Computational modeling suggests that these parallel inhibitory pathways can differentially regulate circuit gain and network stability across different inputs and arousal conditions. Thus, the diversity of disinhibitory GABAergic circuits in the neocortex provides a mechanism for state-dependent flexibility in circuit function.

## Introduction

In the neocortex, GABAergic interneurons play critical roles in shaping neural circuit dynamics underlying perception and the generation of behavior ^1–5^. A large literature has established that cortical interneurons can be classified into a variety of groups based on electrophysiological, anatomical, molecular, and hodological properties ^1,2,6^. For example, fast-spiking interneurons expressing the calcium binding protein parvalbumin (PV-INs) make inhibitory synapses near the peri-somatic regions of excitatory pyramidal neurons and exert well-documented control over the magnitude and timing of spike output ^7–9^. In contrast, regular- or burst-firing interneurons expressing the peptide somatostatin (SST-INs) target pyramidal neuron dendrites and influence synaptic integration and spike output ^10–13^. Other types, such as those expressing vasoactive intestinal peptide (VIP-INs), also innervate pyramidal cell dendrites but may function primarily through their control of other interneurons ^6,14–16^.

Indeed, GABAergic inhibition of interneurons that leads to increased pyramidal cell output via disinhibition has emerged as a key motif supporting cortical circuit dynamics ^6,15,17–20^. In particular, inhibition of SST-INs may play a critical role in the modulation of cortical activity. Anatomical and electrophysiological studies from brain slices documented the innervation of SST-INs by VIP-INs in both prefrontal and visual areas ^6,14,16,21^, while in vivo studies provided evidence that VIP-IN-mediated inhibition of SST-INs occurs and contributes to increased pyramidal cell activity during whisking and locomotion ^15,22–24^. Similarly, long-range projections from frontal to visual areas may enhance sensory responses via VIP-IN-mediated inhibition of SST-INs ^17^. Anatomical and synaptic evidence also indicates that PV-INs directly innervate SST-INs and are capable of evoking postsynaptic inhibition ^21,25^, although the functional role for this pathway in vivo remains unclear. Together, these findings suggest that parallel inhibition of SST-INs in the neocortex might provide diverse mechanisms for circuit control across varying behavioral demands.

Neuromodulation of cellular excitability and synaptic transmission is also a critical mediator of behavioral state-dependent cortical dynamics, with both acetylcholine (ACh) and norepinephrine (NE) demonstrated to play key roles in shaping neural activity during fluctuations in arousal levels ^26–29^. Cholinergic excitation is hypothesized to drive increased activity of VIP-INs during locomotion and resulting cortical disinhibition ^15,30^. In contrast, presynaptic cholinergic receptors can act to suppress GABA release by PV-INs onto target pyramidal neurons, reducing their contribution to feedforward inhibition ^31^. The action of NE on GABAergic circuits is less clear, but early reports provided examples of both enhancement and suppression of synaptic inhibition in the neocortex ^32,33^. The role of neuromodulation in shaping GABAergic inhibition of interneurons is unknown but has the potential to potently influence the role of cortical disinhibition across behavioral states.

Here, we sought to understand both the functional differences in parallel inhibitory inputs to SST-INs within superficial primary visual cortex (V1) and the regulation of these connections by neuromodulators like ACh and NE. Combining optogenetic stimulation and whole-cell recordings, we found that PV-INs innervate proximal somatodendritic compartments of SST-INs and evoke strong, fast synaptic currents. In contrast, VIP-INs target more distal somatodendritic regions and evoke slower currents that distinctly utilize α5 subunit-containing GABA_A_ receptors. Pharmacological manipulation then revealed that activation of muscarinic cholinergic receptors strongly suppresses PV-IN output onto SST-INs, while activation of β-adrenergic receptors facilitates VIP-IN output, suggesting that variation in arousal alters the balance of inhibitory inputs to SST-INs. We explored the functional consequences of this finding via computational modeling. Our results suggest that stronger PV-IN-mediated inhibition of SST-INs, as expected at low arousal levels, increases the gain of the local circuit in response to feedforward input and can enhance overall network stability. Overall, these findings demonstrate that neuromodulation of parallel inhibitory inputs to SST-INs in V1 can enhance the flexibility of circuit dynamics under distinct behavioral conditions.

## Results

In this study, we used the transgenic GIN mouse line, in which enhanced green fluorescent protein (eGFP) is expressed under control of the GAD67 promoter ^34^, to identify and record from somatostatin-expressing interneurons (SST-INs) in the primary visual cortex (V1). Prior studies showed that this line labels 30-40% of SST-INs in layer 2/3 ^35^, although whether they represent a distinct subclass remains unclear. Therefore, we carried out spatial transcriptional profiling of layer 2/3 GIN cells in V1 to characterize expression of several marker genes used to classify SST-INs, including somatostatin (*SST*), calretinin (*CALB2*), cholecystokinin (*CCK*), corticotropin releasing hormone (*CRH*), extracellular leucine-rich repeat and fibronectin type iii domain-containing 1 (*ELFN1*), ETS variant transcription factor 1 (*ETV1*), heparanase (*Hpse*), prodynorphin (*Pdyn*), and the vesicular GABA transporter (VGAT, *Slc32A1*)^36^.

**Figure 1.**
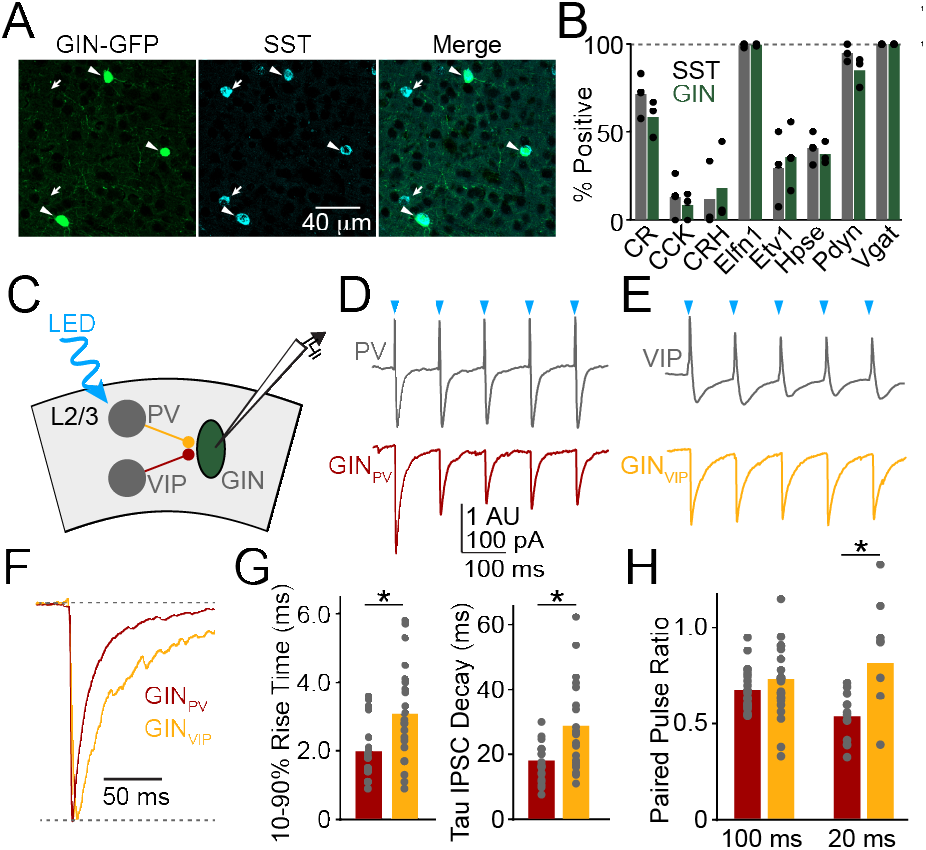
Inhibitory inputs onto L2/3 SST-INs in V1. **A**, Example images showing GFP-expressing GIN cells (right), mRNA labeling for SST (middle), and merge (right). Arrowheads indicate SST-positive GIN cells. Arrows indicate SST-positive non-GIN cells. **B**, Population data showing percentage of all SST-positive cells (gray) or GIN cells (green) expressing each marker. Bars indicate average values, dots indicate individual animals. CR:calretinin, CCK: cholecystokinin, CRH: corticotropin releasing hormone, Elfn1: Extracellular leucine rich repeat and fibronectin type III domain, Etv1: ETA variant transcription factor 1, Hpse: heparanase, Pdyn: prodynorphin, Vgat: Vesicular GABA transporter. **C**, Schematic illustrating experimental setup, with optogenetic stimulation of either PV- or VIP-INs and whole-cell recordings of GIN cells. **D**, Example loose patch recording of PV-IN (top) and whole-cell recording of GIN cell (bottom) showing responses to optical stimulation of PV-INs. **E**, As in (D) for stimulation of VIP-INs. **F**, Example amplitude-normalized PV-IPSC (red) and VIP-IPSC (yellow), illustrating distinct decay kinetics. **G**, Population data showing the 10%-90% rise time (left) and tau of decay (right) for PV-(red) and VIP-IPSCs (yellow). Bars indicate average values and dots represent individual cells. **H**, Population data showing the paired pulse ratios for indicated inter-stimulus intervals for PV-(red) and VIP-IPSCs (yellow). Bars indicate average values and dots represent individual cells. * indicates p<0.05.

Within layer 2/3 of V1, all GIN cells were positive for both VGAT and SST, while ∼50% (74/150) of SST-INs were labeled by GFP (Figure 1A). To determine whether GIN cells make up a distinct subpopulation, we compared transcription expression patterns between them and all SST-positive cells. Results showed high similarity between the two groups (Figure 1B, Supplemental Figure 1), suggesting that at least within layer 2/3 of V1, GIN cells may comprise a broadly representative subset of SST-INs.

We next sought to characterize the functional properties of inhibitory synapses targeting SST-INs in V1, focusing on PV-INs and VIP-INs as candidate presynaptic inputs. We crossed the GIN mouse line to either a PV-Cre ^37^ or VIP-Cre ^38^ line and injected offspring with an adenoassociated viral vector driving Cre-dependent expression of the light-gated excitatory opsin ChroME and a bicistronic red fluorophore mRuby3 ^39^. We then prepared acute slices from adult mice and performed whole-cell recordings from GFP-identified GIN cells (putative SST-INs), using brief light pulses directed through the microscope objective to activate ChroME-expressing PV- or VIP-INs (Figure 1C). Light pulses reliably evoked single spikes in the target presynaptic populations confirmed by loose-patch recordings of mRuby3-positive cells (Fig. 1D-E). Moreover, optical stimulation also produced corresponding inhibitory postsynaptic currents (IPSCs) in recorded GIN cells (Figure 1D-E).

Absolute IPSC magnitudes evoked by widefield optogenetic stimulation are difficult to interpret, given variation in expression levels across slices. However, IPSC kinetics were notably faster for PV-evoked IPSCs (PV-IPSCs) versus VIP-evoked IPSCs (VIP-IPSCs, Figure 1F). Indeed, both the rise time (PV: 2.0 ± 0.1868 ms, n=22 cells vs. VIP: 3.1 ± 0.302 ms, n=22 cells ; p=0.0026, unpaired t-test) and the decay time constant (PV: 17.5 ± 1.187 ms, n=22 cells vs. VIP: 28.2 ± 3.01 ms, n=21 cells; p=0.0017, unpaired t test) were significantly lower for PV-IPSC versus VIP-IPSC data (Figure 1G). We also compared the short-term plasticity of both pathways to repeated optogenetic activation. Both PV- and VIP-inputs exhibited moderate paired-pulse depression at 100 ms and 20 ms intervals that was significantly greater for PV-IPSCs at the shorter interval (for 100 ms, 0.67 ± 0.023, 22 cells vs. 0.74 ± 0.042, 23 cells for PV- vs. VIP-; p=0.2057; for 20 ms, 0.53 ± 0.034, n= 13 cells vs. 0.82 ± 0.104, n= 9 cells for PV- vs. VIP-; p=0.0062, unpaired t test; Figure 1H). Overall, these results confirm that both PV and VIP-INs make robust inhibitory connections with layer 2/3 SST-INs in superficial mouse V1.

Widefield optogenetic stimulation limits interpretation of absolute IPSC amplitudes and may confound synaptic dynamics due to direct excitation of axon terminals. Therefore, we used 2-photon stimulation of ChroME-expressing presynaptic cell bodies to investigate differences between PV- and VIP-inputs to SST-INs. Spiral scanning over individual presynaptic cells reliably evoked short-latency single action potentials with high spatial precision (Supplemental Figure 2). We then used this approach to stimulate either PV- or VIP-INs while recording postsynaptic GIN cells (Figure 2A). We observed a postsynaptic response in 22/68 (32.4%) attempts for PV-IN inputs and 25/64 (39%) attempts for VIP-INs, suggesting similar connection probabilities for the two pathways. Evoked responses under these conditions reflect unitary IPSCs (arising from a single presynaptic neuron), also allowing us to compare amplitudes for the two inputs (Figure 2B). Results showed that unitary IPSCs arising from PV-INs were significantly greater in magnitude than those arising from VIP-INs (PV, 80.2±26.2 pA, n=22 connections; VIP, 23.1±4.6 pA, n=25 connections; p=0.0275, Student’s t-test, Figure 2C) and exhibited significantly faster rise time kinetics (PV, 1.3±0.2 ms, n=22 connections; VIP, 2.5±0.4 ms, n=25 connections; p=0.0179, Student’s t-test, Figure 2C). We again examined short-term plasticity following repeated stimulation, finding that for unitary responses, PV-IPSCs but not VIP-IPSCs showed significant paired-pulse depression (PV, 0.43±0.07, n=22 connections, p<0.0001 Student’s t-test versus 1.0; VIP, 1.28±0.4, n=25 connections, p=0.8677 Student’s t-test versus 1.0; p=0.0306, Student’s t-test PV vs. VIP, Figure 2D).

**Figure 2.**
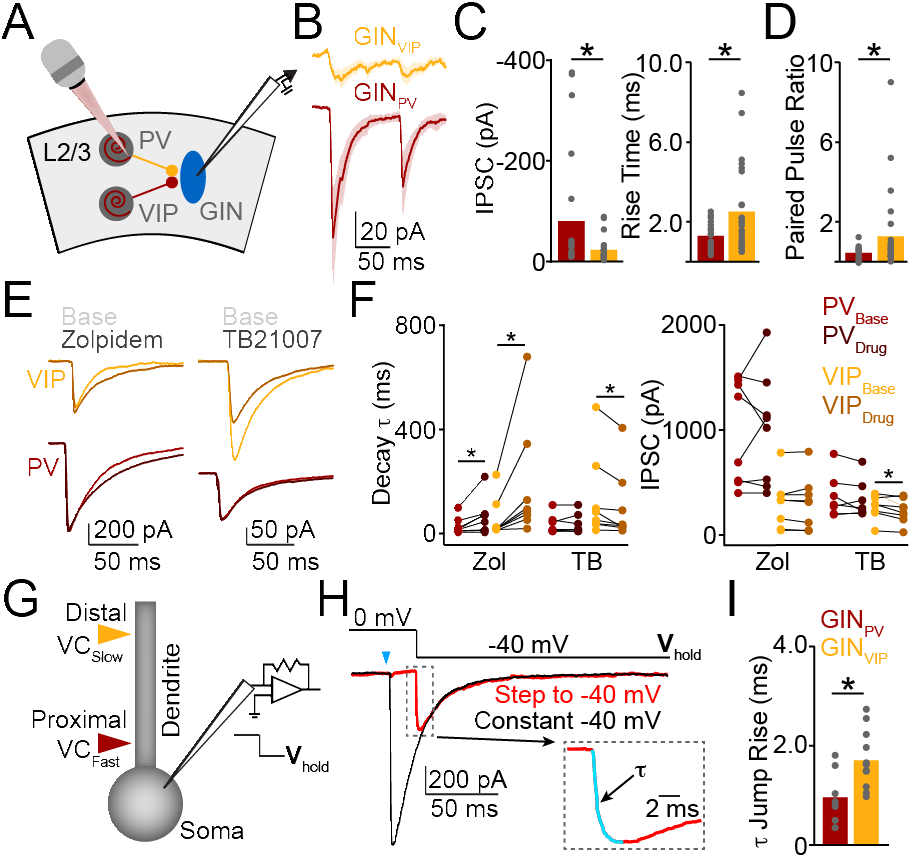
Mechanisms underlying distinct kinetics of PV- and VIP-IN output. **A**, Schematic illustrating experimental setup, with 2-photon optogenetic stimulation targeting individual PV- or VIP-INs. **B**, Example unitary IPSCs evoked in GIN cells by stimulation of PV- (red) or VIP-INs (yellow). **C**, Population data showing IPSC amplitude (left) and rise time (right) for unitary PV-(red) and VIP-IPSCs (yellow). Bars indicate average values and dots represent individual synaptic connections. **D**, Population data showing paired pulse ratio values for unitary PV-(red) or VIP-IPSCs (yellow). Bars indicate average values and dots represent individual synaptic connections. **E**, Example traces showing PV-(red) or VIP-IPSCs (yellow) before (light color) or after (dark color) application of either zolpidem or TB21007. **F**, Population data showing change in IPSC decay (left) and IPSC amplitude (right) for PV-(red) and VIP-IPSCs (yellow) induced by zolpidem or TB21007 (TB). **G**, Schematic illustrating voltage jump protocol. **H**, Example traces indicating voltage-jump protocol. **I**, Population data showing tau of voltage jump-evoked currents for PV- (red) ad VIP-IPSCs (yellow). Bars indicate average and dots represent individual cells. * indicates p<0.05.

Prior studies found that the molecular composition of postsynaptic receptors at synapses formed by different interneuron types onto pyramidal cells can vary, contributing to variation in IPSC kinetics ^40–42^. In particular, receptors containing α5 subunits exhibit substantially longer decay times ^43,44^. Therefore, we compared the contribution of different alpha subunits to the IPSCs measured in SST-INs. Application of the α1 subunit-specific positive modulator zolpidem significantly slowed the decay times for both PV- and VIP-IPSCs (PV-, baseline=30.9±10.0 ms, zolpidem=73.9±25.3 ms, n=8, p=0.0365; VIP-, baseline=49.8±25.4 ms, zolpidem=192.3±80.3, n=8, p=0.0369; paired t test) but did not alter IPSC amplitudes (PV-, baseline=982.5±176.4 pA, zolpidem=1007±190.4, n=8, p=0.8417; VIP-, baseline=320.8±84.3 ms, zolpidem=320.4±89.9, n=8, p=0.9802; paired t test; Figure 2E-F). In contrast, the α5-specific negative modulator TB21007 significantly accelerated the kinetics and decreased the amplitude of VIP-IPSCs (Decay, baseline=138.4±58.4 ms, TB21007=99.6±47.4 ms, p=0.0251; Amplitude, baseline=277.4±38.2, TB21007=226.1±37.7ms, p= 0.0290; paired t test) but did not alter PV-IPSCs (Decay, baseline=48.0±15.9 ms, TB21007=52.0±16.9 ms, n=7, p=0.6006; Amplitude, baseline=335.2±74.9 pA, TB21007= 300.7±64.24 pA, n=7, p= 0.2919; paired t test; Figure 2E-F). These results indicate that α1 subunits are present at both populations of inhibitory postsynaptic contacts, but α5 subunits appear to be selectively localized at synapses formed by VIP-INs.

**Figure 3.**
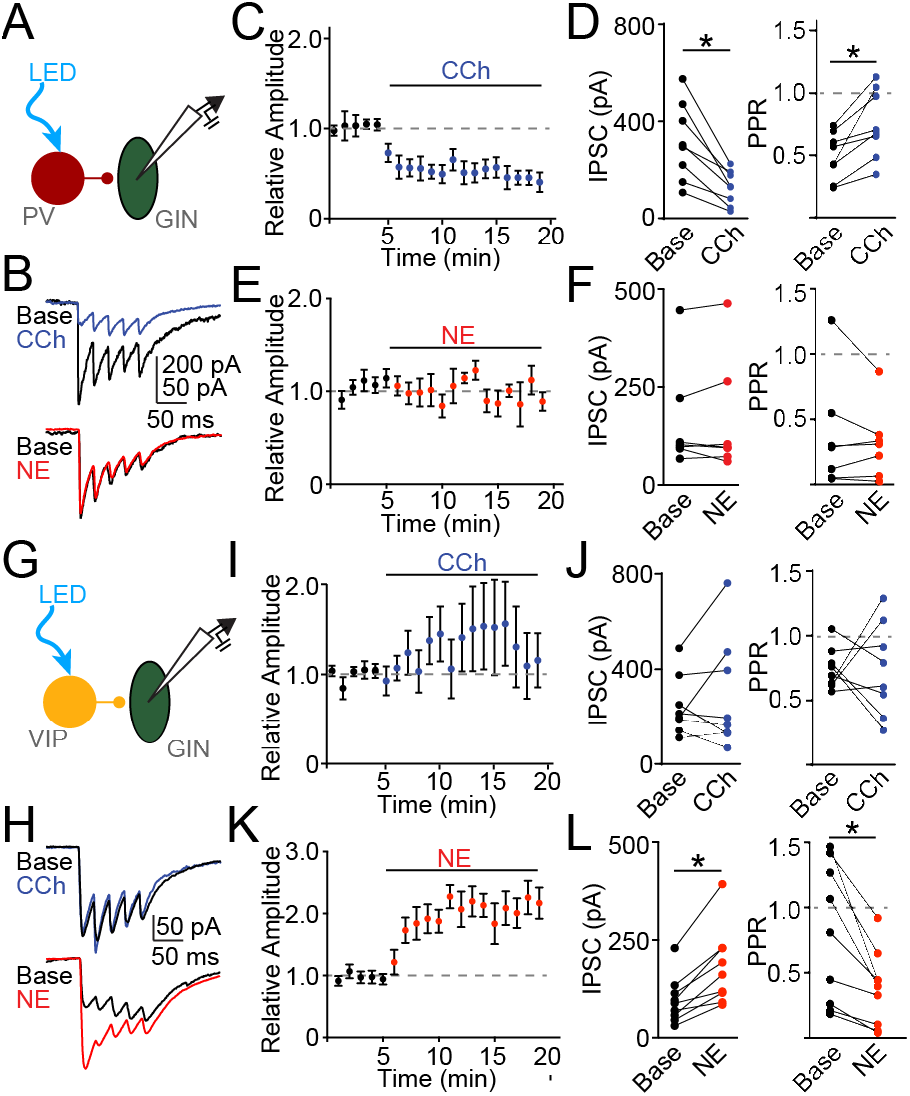
Neuromodulation of PV- and VIP-IPSCs. **A**, Schematic illustrating experimental setup to record PV- IPSCs in GIN cells. **B**, Example PV-IPSCs evoked before or after application of carbachol (CCh, blue) or norepinephrine (NE, red). **C**, Time-course showing average PV-IPSC amplitude during flow-in of CCh. **D**, Population data showing PV-IPSC amplitude (left) and paired-pulse ratio (right) before and after CCh application. Connected dot pairs represent individual cells. **E**, Time-course showing average PV-IPSC amplitude during flow-in of NE. **F**, Population data showing PV-IPSC amplitude (left) and paired-pulse ratio (right) before and after NE application. Connected dot pairs represent individual cells. **G**, Schematic illustrating experimental setup to record VIP-IPSCs in GIN cells. **H**, Example VIP-IPSCs evoked before or after application of carbachol (CCh, blue) or norepinephrine (NE, red). **I-L**, As in (C-F) for VIP-IPSCs. * indicates p<0.05.

GABAergic synapses can also vary in their subcellular localization, with PV-INs primarily targeting perisomatic regions of pyramidal neurons while VIP- and SST-INs primarily target dendritic compartments ^2,11,16^. Such anatomical differences might also extend to synapses formed onto other interneurons and shape measured IPSC kinetics via electrotonic filtering of more distal synaptic currents. To examine this possibility, we used a voltage-jump protocol while recording from postsynaptic GIN cells and optogenetically stimulating PV- or VIP-INs ^45,46^. This method involves holding the cell near the reversal potential for the synaptic current (∼0 mV for GABA_A_ receptors under our experimental conditions) and then stepping the holding potential to a different value (Figure 2G-H). The kinetics of the resulting non-capacitative current are determined by the timing at which the synaptic conductance can be clamped to the new potential, independent from channel activation kinetics. Thus, synaptic inputs more distant from the somatic electrode will be clamped more slowly. We repeated this approach for the two inhibitory inputs, finding significantly slower rise times for voltage jump-evoked currents comparing VIP-versus PV-IPSCs (VIP, 1.7±0.2 ms, n=12; PV: 1.0±0.2, n=8; p=0.0072, unpaired t test, Figure 2I). Overall, our results indicate that VIP-IPSCs exhibit slower kinetics due in part to distinct receptor complements and to more distal localization on the postsynaptic SST-IN.

While disinhibition via suppression of SST-INs is hypothesized to play a critical role in the regulation of cortical activity across behavioral states and fluctuations in arousal levels ^15,17,18^, fluctuations in the release of neuromodulators such as ACh and NE are also implicated in state-dependent cortical dynamics ^20,26,29,47^, potentially through regulation of inhibitory synaptic function. We therefore examined how cholinergic and adrenergic modulation might influence inhibition of SST-INs. We first recorded PV-IPSCs before and after application of the non-selective cholinergic agonist carbachol (Figure 3A-C). Carbachol strongly and significantly suppressed PV-IPSC amplitude while also significantly increasing the paired-pulse ratio (Amplitude, baseline=315.5±56.6 pA, carbachol=126.3±25.1 pA, n= 8, p= 0.0018; Paired pulse ratio, baseline=0.5±0.1, carbachol=0.8±0.1, n=8, p=0.0048; paired t test, Figure 3D), suggesting a presynaptic locus of action. Moreover, the actions of carbachol were completely blocked by pre-application of the specific muscarinic antagonist scopolamine (Supplemental Figure 3). In contrast, application of the broad adrenergic agonist norepinephrine (Figure 3B, E) had no effect on either PV-IPSC amplitude or paired pulse ratio (Amplitude, baseline=162.6±50.9 pA, norepinephrine=165.3±56.0 pA, n=7, p=0.7783; Paired pulse ratio, baseline=0.4±0.12, norepinephrine=0.3±0.1,n= 7, p= 0.4051; paired t test, Figure 3F).

We then performed similar experiments measuring VIP-IPSCs (Figure 3G). Application of carbachol had no significant impact on VIP-IPSC amplitude or paired pulse ratio (Amplitude, baseline=245.3±44.7 pA, carbachol=290.4±83.5 pA, n=8, p=0.4235; paired pulse ratio, baseline=0.7±0.1, carbachol=0.7±0.1, n=8, p=0.9754; paired t test, Figure 3H-J). In contrast, application of norepinephrine significantly increased VIP-IPSC amplitude while decreasing the paired pulse ratio (Amplitude, baseline=96.3±19.9 pA, norepinephrine=179.5±32.4 pA, n=9, p=0.0007; paired pulse ratio, baseline=0.8±0.2, norepinephrine=0.4±0.1, n=9, p=0.004; paired t test, Figure 3H-L), again suggesting a presynaptic locus of action. Moreover, the actions of norepinephrine on VIP-IPSCs were completely blocked by pre-application of the specific β-adrenergic receptor antagonist propranolol and were phenocopied by application of the β agonist isoproterenol (Supplemental Figure 3).

**Figure 4.**
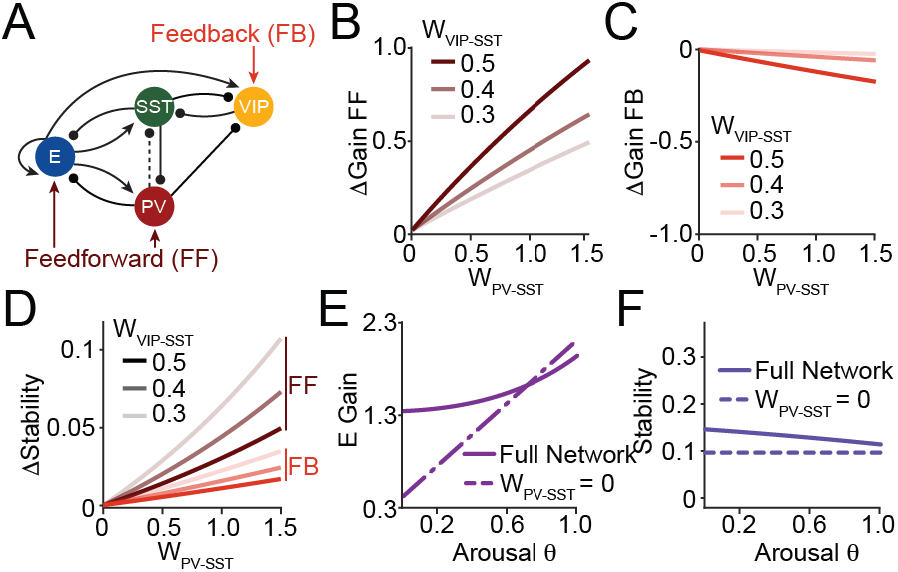
Computational consequences of parallel SST-IN inhibition. **A**, Schematic illustrating the rate-based circuit model including excitatory pyramidal cells (E) and PV-, VIP-, and SST-INs used to explore dynamics under different input (Feedforward versus Feedback) and modulatory regimes. **B**, Gain of circuit response to feedforward (FF) input as a function of increasing PV-IN inhibition of SST-INs. Data are shown for three different strengths of VIP-IN inhibition of SST-INs. **C**, As in (B) for circuit responses to feedback (FB) input. **D**, Network stability as a function of increasing PV-IN inhibition. Data are shown for FF- (brown) and FB- (red) dominated conditions and three different strengths of VIP-IN-mediated inhibition. **E**, Circuit gain for varying levels of arousal (increasing arousal signifies greater proportion of FB versus FF input and stronger VIP-versus PV-IN-mediated inhibition of SST-INs). Dashed line indicates model results with PV-IN inhibition removed. **F**, As in (E) for network stability across arousal levels.

The differential sensitivity of PV- and VIP-IN output to ACh and NE suggest that changes in behavioral state may shift the primary source of SST-IN inhibition in vivo. Prior work also suggests that PV- and VIP-INs may be differentially engaged by feedforward versus feedback projections within the cortex ^17,31,48^. Therefore, we built a computational model to explore whether altering these disinhibitory circuits re-shapes activity in cortical networks. We implemented a recurrent rate-based network including excitatory (E), PV-, VIP-, and SST-IN populations with short-term synaptic depression at the PV-to-SST synapse ^49–51^ (Tsodyks et al., 1998, Wu et al., 2022, Waitzmann et al., 2024). We then applied either feedforward (FF) input to the E and PV populations or feedback (FB) input to the VIP population (see Methods, Figure 4A, Supplemental Figure 4).

We found that when driving the circuit with FF input, increasing the strength of PV-IN-mediated inhibition increased the response gain of the circuit to this input (Figure 4B, Supplemental Figure 4). Moreover, increasing VIP-IN-mediated inhibition further enhanced the gain modulation (Figure 4B, Supplemental Figure 4). In contrast, when the VIP population received FB input, increasing PV-IN-mediated inhibition of SST-INs modestly reduced the response gain to this input (Figure 4C, Supplemental Figure 4), an effect that was again enhanced by stronger VIP-IN-mediated inhibition of SST-INs (Figure 4C, Supplemental Figure 4).

We next examined how inhibition of SST-INs by PV- and VIP-INs influences network stability. Increasing the strength of PV-IN-mediated inhibition of SST-INs generally increased stability, while increasing VIP-IN-mediated inhibition reduced it (Figure 4D, Supplemental Figure 4). The difference in stability between the FF- and FB-dominated conditions reflected differences in the effective strength of PV-to-SST inhibition at steady state due to short-term synaptic depression. Overall, these findings suggest that PV-IN- and VIP-IN-mediated inhibition of SST-INs can work synergistically to control circuit gain for FF versus FB inputs, but with opposing consequences for network stability.

To further compare network dynamics across the transition from low arousal (low ACh and NE, dominant PV-IN-mediated inhibition and FF drive) to high arousal (high ACh and NE, reduced PV-IN-mediated inhibition and FB drive), we modeled a continuous “arousal” parameter θ that interpolates between these conditions (see Methods). We found that, at low arousal values, removal of inhibition by PV-INs drastically reduced both circuit gain and stability (Figure 4E-F), while this manipulation produced minimal impact on the network at high arousal levels. Thus, PV-IN-mediated inhibition of SST-INs plays an important role in maintaining high circuit gain and network stability, particularly during periods of low arousal levels when modulatory influences are reduced and feedforward input provides the dominant source of drive.

## Discussion

In the present study, we found that PV-INs and VIP-INs both provide reliable inhibitory inputs to SST-INs in superficial layers of V1. We identified layer 2/3 SST-INs using the transgenic GIN mouse line, in which GFP is expressed under control of the GAD67 promoter ^34^. Spatial transcriptional profiling showed that GIN cell expression patterns are broadly similar to the overall population of SST-INs, suggesting they do not represent a specific subclass but are generally representative of SST-INs in superficial V1. Thus, in combination with other interneuron-specific labeling strategies (e.g., selective expression of Cre recombinase), the GIN line is a powerful tool for identifying multiple populations in the same mouse ^21^. Nevertheless, prior work has indicated that deeper layer SST-INs comprise a more genetically diverse population ^35,36,52^ and GIN cells are less abundant outside of layer 2/3 ^35^, potentially limiting their utility to superficial cortex.

Our observation that PV-INs make strong inhibitory synapses onto layer 2/3 SST-INs is at variance with early studies that failed to identify this connection ^6^. However more recent work in primary somatosensory cortex also reported inputs to SST-INs from both PV- and VIP-INs ^21,25^. Indeed, our findings from V1 are highly consistent with these reports, showing that the PV-IPSCs are potent and fast, exhibiting high release probability and short-term depression. In addition, our experiments using 2-photon stimulation of individual PV-INs showed similarly strong, depressing responses, indicating that the functional properties observed for this connection are not artifacts arising from bulk optogenetic activation of multiple presynaptic axons.

Our novel experimental results now provide a multifactorial explanation for the relatively faster kinetics of PV-versus VIP-IPSCs onto SST-INs. First, voltage-jump experiments showed that the synapses formed by PV-INs are more proximally located on the somatodendritic membrane versus the more distal synapses formed by VIP-INs, producing stronger electrotonic filtering for the latter. These results echo the well-established perisomatic innervation pattern of PV-INs onto pyramidal neurons as well as observations that VIP-INs target pyramidal dendrites ^1,16^. The mechanisms governing these structural innervation patterns onto pyramidal neurons are only partially understood ^53^. Nevertheless, our results suggest that selective synaptic localization may be a common feature of GABAergic inputs across diverse postsynaptic targets. Second, we found that PV- and VIP-IPSCs are mediated by distinct complements of GABA receptors, with synapses formed by VIP-INs including α5 subunit-containing receptors. This class of receptor is known to exhibit slower kinetics ^43,44^, likely also contributing to the slower VIP-IPSCs. Interestingly, we recently showed that SST-IN inputs to pyramidal cell dendrites also express α5 subunit-containing receptors ^54^, suggesting distal localization of this subunit may be a general principle across cortical cell types. Dendritic α5-containing receptors in pyramidal neurons can also mediate a tonic GABAergic current that paradoxically enhances dendritic calcium signaling in response to back-propagating action potentials ^54^. The dendrites of SST-INs are known to exhibit calcium influx in response to the spread of action potential-mediated depolarization ^55^. Thus, activation of α5-receptors by VIP-IN inputs may be well-positioned to mediate tonic currents in SST-INs, a hypothesis that remains to be explored by future work.

Neuromodulation by ACh and NE plays a critical role in shaping cortical circuit dynamics, particularly during transitions between behavioral states and arousal levels ^26–29^, while also shaping sensory responses in V1 and visual perceptual thresholds ^20,56,57^. Prior studies have generally focused on the ability of these modulators to directly influence the excitability of excitatory and inhibitory neurons. For example, cholinergic activation of VIP-INs is hypothesized to inhibit SST-IN output during locomotion, enhancing pyramidal neuron firing via disinhibition ^15^. However, relatively fewer studies have examined the consequences of modulation for GABAergic synaptic output. Here, we observed that cholinergic signaling robustly suppresses PV-IPSCs in SST-INs. Our data further suggest that this effect is mediated by presynaptic muscarinic receptors, similar to prior results showing muscarinic suppression of PV-IPSCs onto pyramidal neurons in somatosensory cortex ^31^. Complementary to the actions of ACh, we also found that noradrenergic signaling potentiates the output of VIP-INs, an effect mediated by presynaptic β-type adrenergic receptors. Earlier work provided conflicting reports on the consequences of adrenergic modulation for GABAergic transmission in the cortex ^32,33^. However, to our knowledge, this is the first demonstration that VIP-IN output is directly under neuromodulatory influence.

In vivo, fluctuations in both ACh and NE release across the cortex are linked to changes between behavioral states or arousal levels ^28,29,58^. Our results suggest that transitions to higher arousal levels would involve a shift from PV-IN to VIP-IN-mediated inhibition of SST-INs. Several studies have highlighted cortical disinhibition following suppression of SST-INs as a central motif of cortical circuit function ^14,15,22,23^. Based on our findings, we propose that parallel inhibitory pathways support this operation and that changes in behavioral state and modulatory signaling will determine the relative contribution of PV- and VIP-INs to disinhibition. Our computational approach allowed us to explore the functional consequences of these parallel inhibitory pathways for circuit gain and network stability, reaching an intriguing set of hypotheses. PV-INs are often closely associated with the control of feedforward input carrying afferent information from the external environment to sensory cortex ^1,8,48^. In contrast, VIP-INs are suggested to participate in feedback, top-down control of circuits that provide contextual and executive control of sensory processing ^17^. Our modeling work suggests that the relative contributions of PV- to VIP-mediated inhibition of SST-INs can support high circuit gain across feedforward and feedback operating regimes, while their effects on network stability depend on the underlying connectivity.

In conclusion, we demonstrate the existence of parallel inhibitory pathways targeting SST-INs in V1 with distinct physiological properties and sensitivity to neuromodulation associated with changes in behavioral state. Our modeling results suggest that these pathways can differentially regulate circuit gain and network stability across a range of input and connectivity conditions. Thus, the diversity of GABAergic inhibition of SST-INs represents a novel mechanism by which cortical circuits can be flexibly tuned to match ongoing behavioral needs.

## Supplemental Figures

**Supplemental Figure 1.**
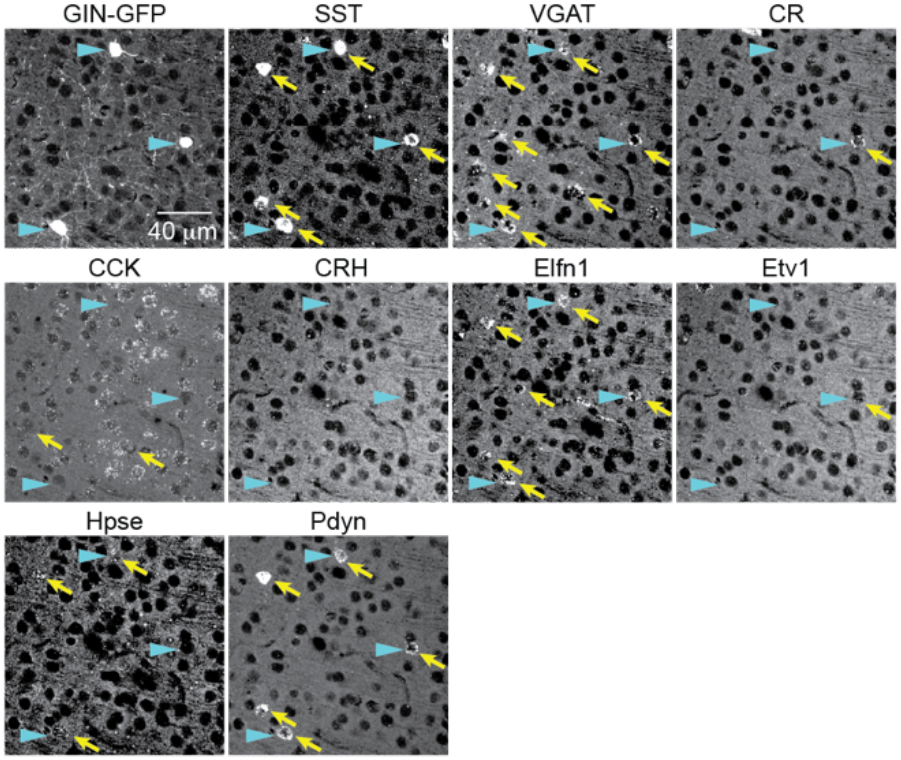
Example images showing RNAscope labeling of GIN cells in layer 2/3 of V1. Panels show data from the same field of view for GIN-GFP, somatostatin (SST), vesicular GABA transporter (VGAT), calretinin (CR), cholecystokinin (CCK), corticotropin releasing hormone (CRH), extracellular leucine rich repeat and fibronectin type III domain (Elfn1), ETA variant transcription factor 1 (Etv1), heparanase (Hpse), and prodynorphin (Pdyn). Blue arrowheads indicate the same GIN cells in each panel. Yellow arrows indicate cells positive for the indicated gene.

**Supplemental Figure 2.**
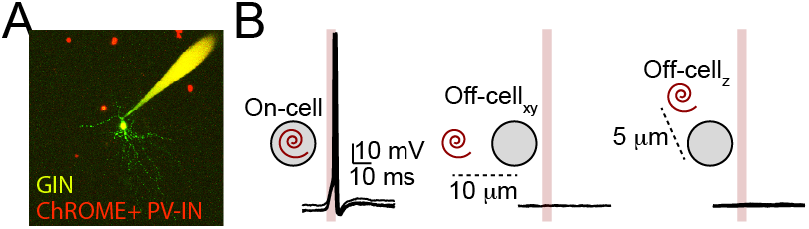
Targeted 2-photon optogenetic stimulation. **A**, Example image showing ChroME;mRuby3-expressing PV-INs (red) and the Alexa Fluor-488-filled patch pipette and GIN cell (green). **B**, Example recordings from ChroME-expressing cell showing reliable action potentials evoked by spiral scanning directly on the cell body (left) but not offset in the x (middle) or z (right) axes.

**Supplemental Figure 3.**
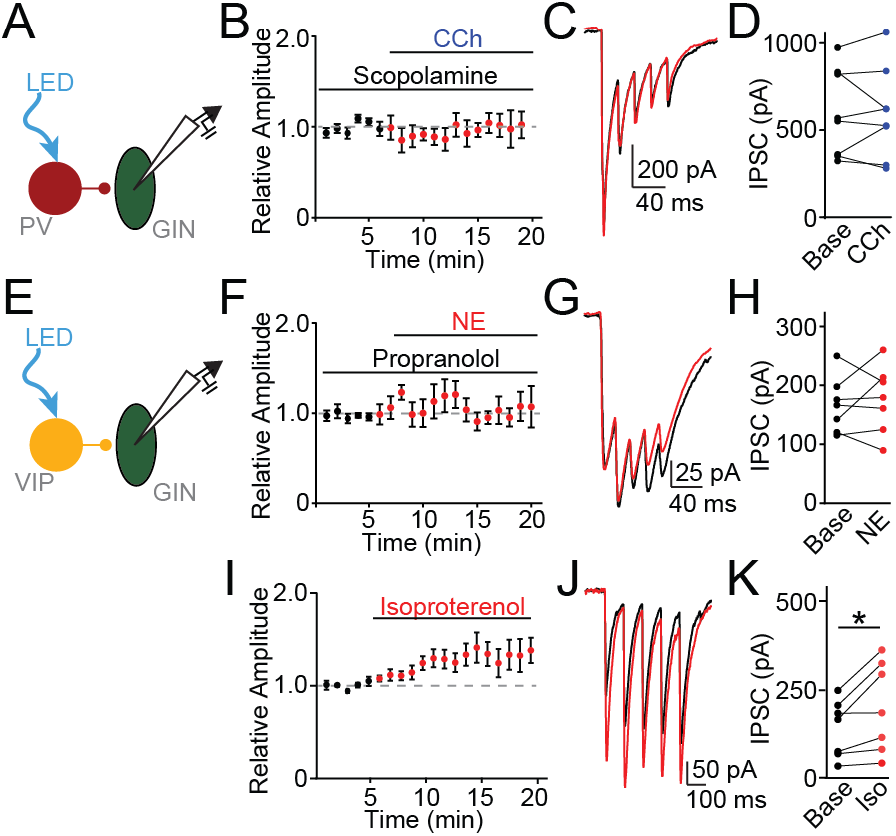
Receptor subtypes underlying neuromodulation of PV- and VIP-IPSCs. **A**, Schematic illustrating experimental setup to record PV-IPSCs in GIN cells. **B**, Time-course showing average PV-IPSC amplitude during flow-in of CCh in the presence of pre-applied scopolamine. **C**, Example PV-IPSCs evoked before or after application of carbachol (CCh, blue) in the presence of pre-applied scopolamine. **D**, Population data showing PV-IPSC amplitude before and after CCh application in the presence of scopolamine. Connected dot pairs represent individual cells. **E**, Schematic illustrating experimental setup to record VIP-IPSCs in GIN cells. **F**, Time-course showing average VIP-IPSC amplitude during flow-in of NE in the presence of pre-applied propranolol. **G**, Example VIP-IPSCs evoked before or after application of NE in the presence of pre-applied propranolol. **H**, Population data showing VIP-IPSC amplitude before and after NE application in the presence of propranolol. Connected dot pairs represent individual cells. **I**, Time-course showing average VIP-IPSC amplitude during flow-in of isoproterenol. **J**, Example VIP-IPSCs evoked before or after application of isoproterenol. **K**, Population data showing VIP-IPSC amplitude before and after isoproterenol application. Connected dot pairs represent individual cells. * indicates p<0.05.

**Supplemental Figure 4.**
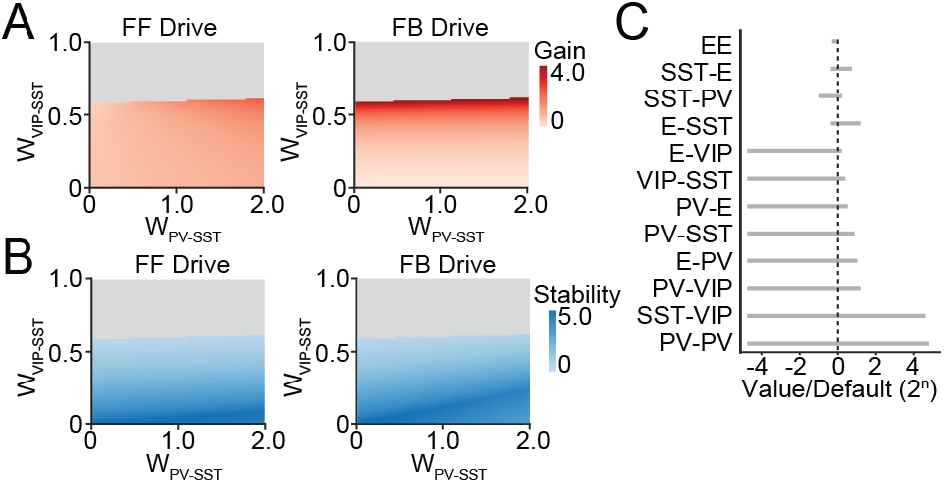
Sensitivity of model circuit to varying inhibition of SST-INs. **A**, Circuit gain in response to feedforward (FF, left) or feedback (FB, right) drive under varied strength of VIP-IN or PV-IN-mediated inhibition of SST-INs. **B**, As in (A) for network stability. **C**, Ranges of network connection weights for which the model yielded stable results.

## Resource Availability

### Lead Contact

Michael J. Higley

## Materials Availability

Requests for reagents used in this study may be made to the Lead Contact.

## Data and Code Availability

All data are presented in the paper and supplementary materials. Any additional information required to reanalyze the data reported in this work paper is available from the Lead Contact upon request (https://github.com/comp-neural-circuits/parallel-inhibition-SST).

## Acknowledgements

The authors wish to thank members of the Higley laboratory and Dr. Jessica A. Cardin for helpful comments during the preparation of this manuscript.

## Authors contributions

Experiments were conceived and designed by JGu and MJH. Experimental data were acquired by JGu and CQC. Modeling data were generated by FV and JGj. Analyses were carried out by JGu, CQC, and FV. Work was directed and supervised by JGj and MJH. Manuscript was written by JGu and MJH and revised by all authors.

## Experimental Methods

### Experimental model

All animal handling was performed in accordance with the guidelines of the Yale Institutional Animal Care and Use Committee. Mice were kept on a 12-hour light-dark cycle in the Yale University School of Medicine animal facilities and provided with food and water *ad libitum*. To generate experimental mice, GIN mice (hemizygotes; RRID:IMSR_JAX:003718) were crossed with interneuron-specific Cre-driver lines (PV-Cre; RRID:IMSR_JAX:008069 and VIP-Cre). Both male and female PV-Cre;GIN and VIP-Cre;GIN mice aged P60-100 days were used for experiments.

### AAV virus injection

To optically stimulate either PV- or VIP-INs, we conditionally expressed the light-activated cationic channel ChroME ^39^ in PV-Cre;GIN or VIP-Cre;GIN mice. Briefly, mice were anesthetized with 1-2% isoflurane and maintained at 37°C. A small burr hole was made through the skull followed by bilateral injection into superficial V1 with 700nl of AAV 2/9-CAG-DIO-ChroME-ST-P2A-H2B-mRuby3 (Addgene 108912) at 3.5mm posterior ad 2.5mm lateral to bregma. Acute slices were prepared at least two weeks following injection.

### Slice preparation

Acute slices of V1 were prepared from PV-Cre;GIN and VIP-Cre;GIN mice (P40-100). Briefly, isoflurane-anesthetized mice were decapitated and coronal slices (300 μm thick) were cut in ice-cold external solution containing (in mM): 100 choline chloride, 25 NaHCO_3_, 1.25 NaH_2_PO_4_, 2.5 KCl, 7 MgCl_2_, 0.5 CaCl_2_, 15 glucose, 11.6 sodium ascorbate and 3.1 sodium pyruvate, bubbled with 95% O_2_ and 5% CO_2_. Slices were then transferred to warm artificial cerebrospinal fluid (ACSF) containing (in mM): 127 NaCl, 25 NaHCO3, 1.25 NaH2PO4, 2.5 KCl, 1 MgCl2, 1.2 CaCl2 and 15 glucose, bubbled with 95% O2 and 5% CO2. After incubating at 34°C for 30 minutes, the slices were maintained at room temperature for at least 30 min before use.

### Electrophysiology

Experiments were conducted at 32°C in a submersion-type recording chamber. Whole-cell voltage-clamp recordings were obtained from GFP-expressing cells (identified via epifluorescence) using infrared differential interference contrast. Cells were located within layer 2/3, 200-300 μm from the pial surface. To record inhibitory currents at a holding potential of −70 mV, glass electrodes (3.0-3.5 MΩ) were back filled with a high chloride internal solution containing (in mM): 100 CsCl, 30 CsGluconate, 10 HEPES, 4 MgCl_2_, 4 Na_2_ATP, 0.4 NaGTP and 10 sodium creatine phosphate, adjusted to pH 7.3 with CsOH. For all experiments, the series resistance was kept <25 MΩ, and experiments were discarded if this value changed more than 20% during the recording. Electrophysiological recordings were made using a Multiclamp 700B amplifier (Molecular Devices), filtered at 4 kHz, and digitized at 10 kHz using ScanImage acquisition software ^59^.

### Optogenetic stimulation

For widefield optogenetic stimulation of PV- and VIP-INs, a 473nm laser (Optoengine) was directed through the microscope objective. Brief pulses (1 ms, 2 mW at the sample) of light reliably evoked postsynaptic currents. For 2-photon optogenetic stimulation…

### Pharmacology

For all experiments, the recording ACSF included 10 μM NBQX to block AMPA receptors and 10 μM CPP to block NMDA receptors. In a subset of experiments, the following compounds were added to the recording ACSF (in μM): 1 zolpidem, 0.1 TB21007, 20 carbachol, 10 scopolamine, 10 norepinephrine bitartrate, 1 propranolol, 1 isoproteranol. All compounds were purchased from Tocris. All compounds were dissolved in water except TB21007 and NBQX in DMSO.

### RNAscope Assay

Formalin-fixed paraffin-embedded (FFPE) tissue was used for RNAscope. For preparation of FFPE tissue, GIN mice were perfused with 4% paraformaldehyde, and brains were post-fixed in 4% PFA, followed by dehydration using tertiary-butanol and tissue clearing using Histochoice. The brains were then infiltrated with and embedded in paraffin. 5-micron thick sections were prepared using a microtome. *In situ* detection of CALB2, CNR, CCK, CRH, ELFN1, ELFN2, ETV1, HPSE, PDYN, SST, and VGAT was performed using the RNAscope Hiplex V2 assay from ACD Bio, following the manufacturer’s protocol. Sections were baked at 60°C for 1 hour, followed by deparaffinization with Histochoice and tertiary butanol. The first round of imaging was conducted after this step, using a confocal microscope. Fields of view containing GFP-expressing cells in the V1 were imaged, and coordinates were recorded for subsequent imaging. The sections then underwent probe hybridization, amplification, and fluorophore labeling, followed by imaging of the previously recorded fields of view. The DAPI imaging was used to verify that the same fields of view were imaged. The fluorophores were then cleaved, and three additional rounds of labeling and imaging were performed in the previously imaged fields of view. To quantify the expression of the various probes across the cells, images from multiple rounds were registered using ACD Bio’s HiPlex Image Registration program to generate 12-channel images containing all the probes and DAPI. Analysis was carried out using QuPath, and RNAscope signal was quantified as the number of spots per cell using QuPath’s subcellular detection function. The minimum spot size was set to 0.5 um2, and the maximum to 2 um2. Spots exceeding this size were detected as clusters and the number of spots per cluster were calculated using an expected spot size of 1 um2.

### Quantification and Statistical Analysis

Offline inspection and analysis of electrophysiological recordings was performed using IgorPro (Wavemetrics). IPSC amplitudes were calculated by finding the peak of the current traces. Statistical comparisons were carried out as described in the text and performed using Prism.

### Network computational model

We modeled a cortical microcircuit of four interacting populations: pyramidal cells (E), PV, SST and VIP interneurons. The rate dynamics follow the following equations (Waitzmann et al., 2024):

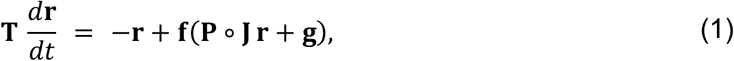

where **T** = diag(*τ_E_*, *τ_P_*, *τ_S_*, *τ_V_*) with *τ_X_* = 10 ms, **r** = (*r_E_*, *r_P_*, *r_S_*, *r_V_*)^T^ is the vector of population rates, **f** a vector of rectified-linear input-output functions, **J** the connectivity matrix, **P** the matrix of STP variables (**P***_i_*_j_ = 1 on synapses without STP), ∘ the element-wise product, and **g** the external inputs. The connectivity matrix is defined as:

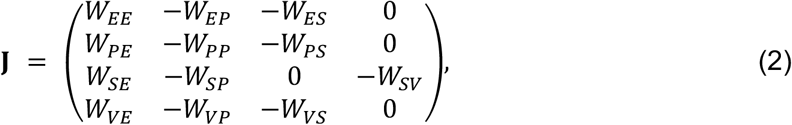

whose structural connectivity follows the known connectivity of these cell classes (Pfeffer et al., 2013, see Table S1). Synaptic signs are included in **J**, so *W_X_*_F_ = |*J_X_*_F_|. The background inputs **g** are chosen such that the rate set-point **r**^∗^ = (3,5,2,2) Hz is a fixed point of the full system. The circuit is in the inhibition-stabilized regime (ISN) (*W_EE_* > 1: the E subnetwork alone is unstable and is stabilized by feedback inhibition).

The PV-to-SST connection expresses short-term depression, modeled with Tsodyks–Markram dynamics through a resource variable *x_SP_* (Tsodyks et al., 1998; Waitzmann et al., 2024):

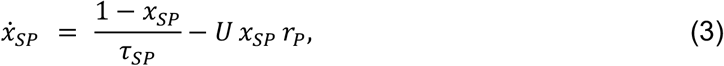

with *τ_SP_* = 100 ms. The steady-state STP variable at PV rate *r_P_* is

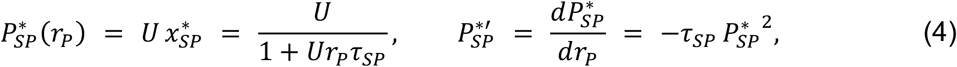

with release probability *U*; all other entries of **P** equal 1 (and 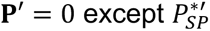). The parameter *U* sets both the first-pulse amplitude (∝ *UW_SP_*) and the paired-pulse ratio 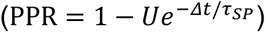.

We modeled the VIP-to-SST synapse as a static synapse, focusing on its steady-state efficacy rather than its transient synaptic dynamics. We therefore did not include short-term plasticity at this connection. The effects of varying VIP-to-SST inhibitory strength on circuit gain and stability were examined separately.

We defined as “arousal” a single parameter *θ* ∈ [0,1] acting on two elements: (1) cholinergic modulation of the PV-to-SST release probability, *U*(*θ*) = 0.5 − 0.35 *θ*, interpolating between the measured control (*U_low_* = 0.5) and carbachol (*U_high_* = 0.15) states; and (2) input routing, *I*(*θ*) = (1 − *θ*) ⋅ FF + *θ* ⋅ FB, where FF targets E and PV and FB targets VIP. Population firing rates were allowed to vary with *θ*, rather than being constrained to remain at the reference operating point.

We linearized Eq. (1) at the fixed point **r**^∗^, replacing the STP variables by their steady-state values (Waitzmann et al. 2024):

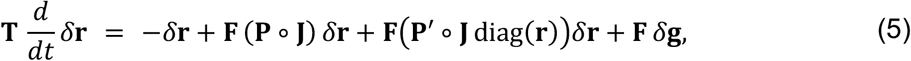

where **F** is the diagonal matrix of input–output slopes at the fixed point (**F** = **1**, the identity, for rectified-linear units with all populations active), **P**′ contains the derivatives of the STP variables with respect to the presynaptic rate, and *δ***g** the input perturbations. The steady-state solution defines the response matrix

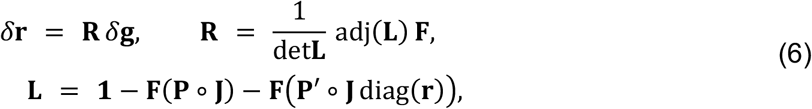

where 1 denotes the identity matrix; the element **R***_i_*_j_ gives the steady-state rate change of population *i* per unit input perturbation to population *j*. The channel gains of the E population are *g*_FF_ = **R***_EE_* + **R***_EP_* = [adj(*L_EE_*) + adj(*L_EP_*)\/det(**L**) (equivalent to feedforward input FF to E and PV) and *g*_FB_ = **R***_EV_* = adj(*L_EV_*)/det(**L**) (equivalent to feedback input FB to VIP). Static (no-STP) controls are the same circuit evaluated at *k* = *W_SP_* (**P** = 1, i.e., *U* = 1).

The PV-to-SST connection enters the linear response through a single number, the effective coupling *k* = *c_SP_W_SP_*. The two gains, under FF and FB, can be rewritten as functions of *k*:

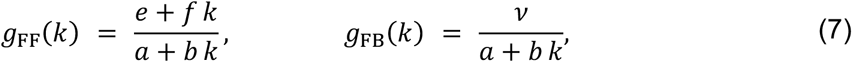

where *a* = det**L**|*_k_*_:0_ is the *full* 4 × 4 determinant of the ablated circuit (*W_SP_* = 0, all other connections intact), *e* and *ν* are the corresponding full FF and FB numerators respectively, so that *e*/*a* = *g*_FF_(0) and *ν*/*a* = *g*_FB_(0) are the ablation gains. *b*, *f* and *ν* are defined as:

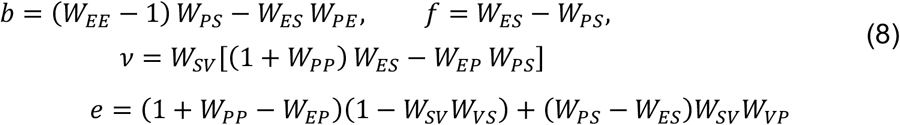

Subtracting the *k* = 0 value from Eq. (7), we obtain

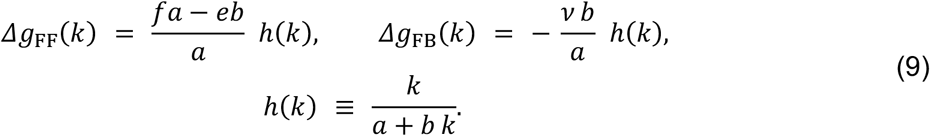

The shape is determined by *a* and *b* since ℎ′(*k*) = *a*/(*a* + *bk*)^2^ > 0 on the stable branch, so ℎ is strictly increasing and monotonic. STP does not change ℎ, but only sets a correction on a given *k* (e.g., *k* = *c_SP_W_SP_* with depression versus *k* = *W_SP_* without).

For the FF gain relative to the ablation (*k* = 0),

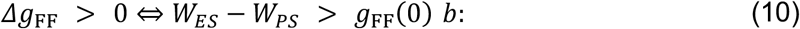

Our result in panel Fig. 4B requires the weights to satisfy

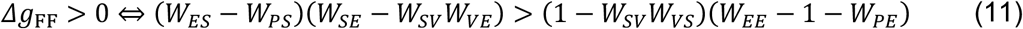

In contrast, the STP-versus-static comparison of the FB gain for the same weight collapses onto *b* alone: both points lie on the same hyperbola, so

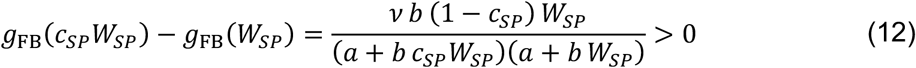

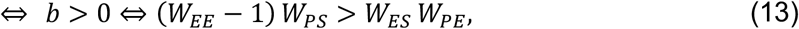

every factor is positive (assuming positive gain *ν* > 0), so the entire comparison reduces to the sign of *b*. Our result in panel E then requires the weights to satisfy (*W_EE_* − 1) *W_PS_* > *W_ES_ W_PE_*. To respect both conditions at the same time, we required:

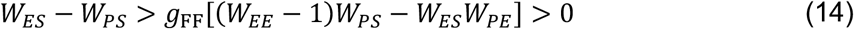

Network stability is quantified as −max Re *λ*, the sign-flipped real part of the leading eigenvalue of the rate Jacobian −**T**^-1^**L** evaluated at the operating point (with the depressing synapse at its steady-state efficacy), so that positive values indicate a stable fixed point and larger values a faster decay of perturbations. This analysis does not explicitly include the transient dynamics of short-term synaptic plasticity. Unlike the gains, the stability matrix has no closed form. Therefore, we evaluated it on a grid of 26,214,400 ISN weight configurations (*W_EE_* ∈ {1.1,1.3,1.5,1.8}, *W_SP_*, *W_SV_* ∈ {0.1,0.7,1.2,1.8}, while the remaining weights each taking the values ∈ {0.1,0.7,1.2,1.8}). We found that the direction of the stability change *ΔStability* divides the parameter space into two comparable regimes: stability declines monotonically with arousal in ≈ 57% of sampled configurations and increases in ≈ 43% (including configurations with a local stability maximum). The circuit shown belongs to the rising class, consistent with previous results (Bos et al., 2025). However, other connectivity configurations exhibited decreasing stability, indicating that the relationship between arousal and network stability depends on the underlying connectivity.

When modelling arousal-dependent changes (Figure 4E-F), we evaluated the network at steady state for each value *θ*. The differences in stability in Figure 4F depend on changes in the efficacy of *W_SP_*, and not on the changes in input strength. Furthermore, the intersection of the gain curves in Figure 4E also depends on the strength of *W_SV_* inhibition. Changing this connection can shift the arousal level at which the curves intersect or eliminate the intersection altogether for certain values of *W_SV_*.

**Table S1:** Model parameters.

| Parameter | Description | Value |
| --- | --- | --- |
| $\tau_E, \tau_P, \tau_S, \tau_V$ | Rate time constants | 10 ms |
| $\tau_{SP}$ | PV-to-SST depression recovery | 100 ms |
| $U_{low}$ | Release probability, control | 0.5 |
| $U_{high}$ | Release probability, carbachol | 0.15 |
| $\mathbf{g}_{FF}$ | FF input: unit drive to E and PV | (1,1,0,0) |
| $\mathbf{g}_{FB}$ | FB input: unit drive to VIP | (0,0,0,1) |
| $W_{EE}$ | E→E recurrent excitation | 1.3 |
| $W_{EP}$ | PV→E inhibition | 0.7 |
| $W_{ES}$ | SST→E inhibition | 1.2 |
| $W_{PE}$ | E→PV excitation | 0.1 |
| $W_{PP}$ | PV→PV inhibition | 0.1 |
| $W_{PS}$ | SST→PV inhibition | 0.7 |
| $W_{SE}$ | E→SST excitation | 1.2 |
| $W_{SP}$ | PV→SST inhibition | 1.5 |
| $W_{SV}$ | VIP→SST inhibition | 0.5 |
| $W_{VE}$ | E→VIP excitation | 1.2 |
| $W_{VP}$ | PV→VIP inhibition | 1.2 |
| $W_{VS}$ | SST→VIP inhibition | 0.1 |

